# Unveiling the evolutionary history of lingonberry (*Vaccinium vitis-idaea* L.) through genome sequencing and assembly of European and North American subspecies

**DOI:** 10.1101/2023.10.19.563133

**Authors:** Kaede Hirabayashi, Samir C. Debnath, Gregory L. Owens

## Abstract

Lingonberry (*Vaccinium vitis-idaea* L.) produces tiny red berries that are tart and nutty in flavour. It grows widely in the circumpolar region, including Scandinavia, northern parts of Eurasia, Alaska, and Canada. Although cultivation is currently limited, the plant has a long history of cultural use among indigenous communities. Given its potential as a food source, genomic resources for lingonberry are significantly lacking. To advance genomic knowledge, the genomes for two subspecies of lingonberry (*V. vitis-idaea* ssp. *minus* and ssp. *vitis-idaea* var. ‘Red Candy’) were sequenced and *de novo* assembled into contig-level assemblies. The assemblies were scaffolded using the bilberry genome (*V. myrtillus*) to generate chromosome-anchored reference genome consisting of 12 chromosomes each with total length 548.07 Mbp (contig N50 = 1.17 Mbp, BUSCO (C%) = 96.5%) for ssp. *vitis-idaea*, and 518.70 Mbp (contig N50 = 1.40 Mbp, BUSCO (C%) = 96.9%) for ssp. *minus*. RNA sequencing based gene annotation identified 27,243 genes on the ssp. *vitis-idaea* assembly, and transposable element detection methods found that 45.82% of the genome was repeats. Phylogenetic analysis confirmed that lingonberry is most closely related to bilberry and is more closely related to blueberries than cranberries. Estimates of past effective population size suggested a continuous decline over the past 1–3 MYA, possibly due to the impacts of repeated glacial cycles during Pleistocene leading to frequent population fragmentation. The genomic resource created in this study can be used to identify industry relevant genes (e.g., flavonoid genes), infer phylogeny, and call sequence-level variants (e.g., SNPs) in future research.

## Introduction

*Vaccinium vitis-idaea* L., commonly known as lingonberry, also called partridgeberry or mountain cranberry, is an evergreen dwarf shrub that has cultural, economic, and ecological importance (Debnath and Arigundam 2020). The bright-red coloured berries have been consumed among Indigenous communities in northern North America and Scandinavia as a relish and served with meat or fish in traditional meals (Moerman 2010; Vaara *et al*. 2013). Berry picking has been a cherished cultural practice and nowadays people commonly preserve berries as jams which are becoming more readily available commercially (“Arctic Lingonberry” 2022). A growing body of research suggests that lingonberry fruits and leaves have medicinal benefits to human health such as anticancer, cardioprotective, and neuroprotective properties (reviewed in Ferlemi and Lamari 2016; Kowalska 2021). Despite a long history of utilization as a culturally important food source and its recognized health benefits, the domestication of lingonberry is at its infancy in North America.

Being an evergreen boreal forest understory species, lingonberry propagates vegetatively by forming mat-like clonal communities through rhizomes (Hjalmarsson and Ortiz 1998), or sexually through seeds which are primarily insect pollinated (Jacquemart and Thompson 1996). The species has two recognized subspecies (ssp.) based on their geographical origin: *V. vitis-idaea* ssp. *minus* and ssp. *vitis-idaea* and the species is widely distributed in the circumpolar region (Debnath and Arigundam 2020; Figure 1a). The European subspecies, ssp. *vitis-idaea*, currently has active breeding programs with more than a dozen of cultivars available for commercial production, with improved yield and berry size (Penhallegon 2009; Figure 1b). The North American ssp. *minus*, on the other hand, is considered a wild plant with little breeding efforts taken place. The two subspecies are distinguishable based on several morphological differences as well as genetic differences (Garkava-Gustavsson *et al*. 2005; Debnath 2007; Debnath and Arigundam 2020; Figure 1c). The extent of genomic differences between the two subspecies has not been studied before, and it is somewhat unclear whether they occur sympatrically in the overlapping ranges.

**Figure 1:**
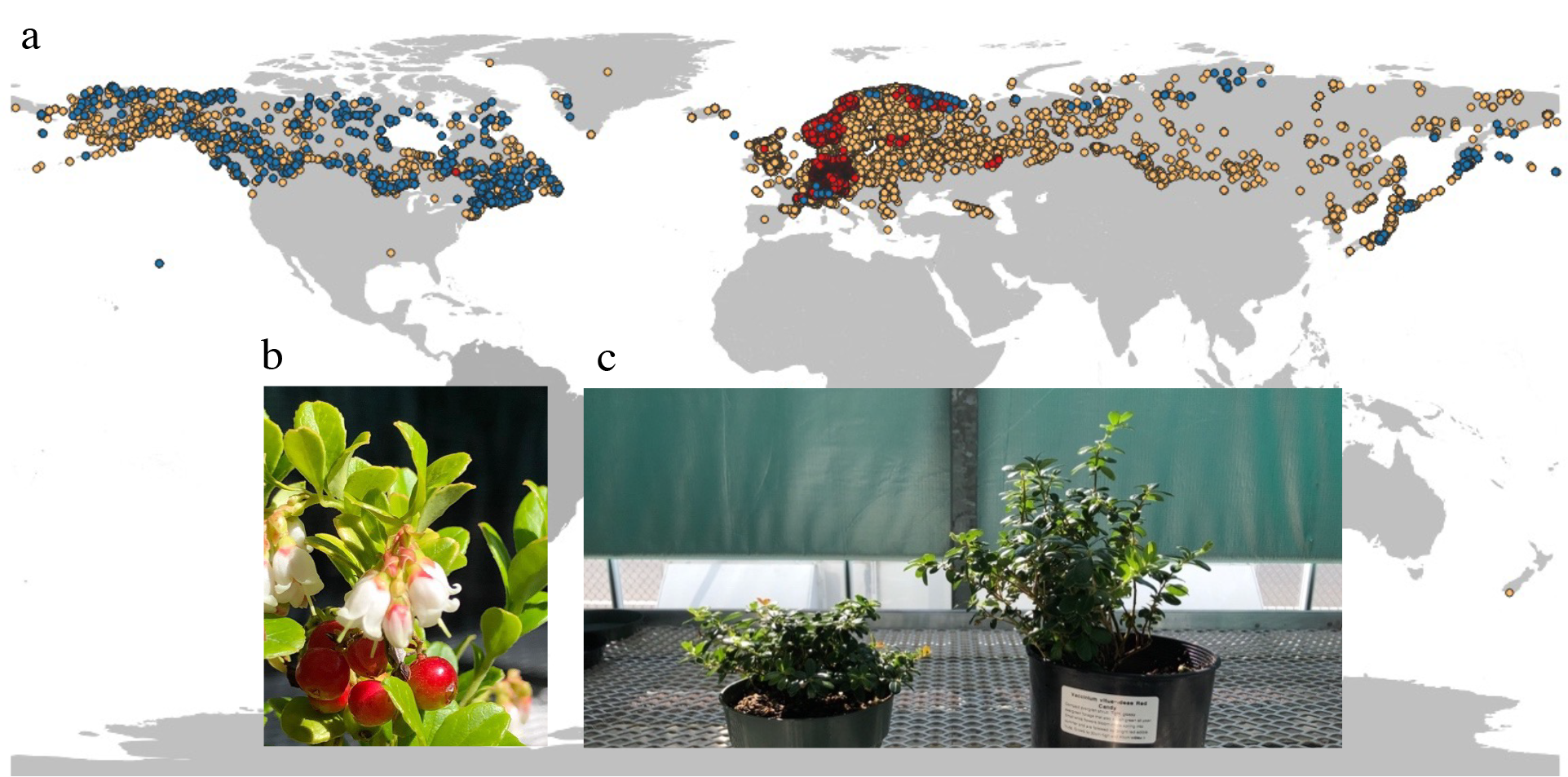
**a)** Worldwide distribution of *Vaccinium vitis-idaea* L. (www.gbif.org) Dots represent occurrence records registered as: *V. vitis-idaea* ssp. *minus* (blue), *V. vitis-idaea* ssp. *vitis-idaea* (red), *V. vitis-idaea* L. ssp. unidentified (yellow). **b)** *V. vitis-idaea* ssp. *vitis-idaea* flowers and fruits. **c)** *V. vitis-idaea* ssp. *minus* (left) and ssp. *vitis-idaea* var. ‘Red Candy’ (right) grown in the greenhouse.

Long-read sequencing technology has fueled exponential growth in the assembly of plant genomes (https://www.plabipd.de/timeline1_view.html); there are at least 1,205 unique flowering plant species genomes assembled at higher than scaffold-level (NCBI search terms: “Magnoliopsida (flowering plants)” “scaffold+”, by May 24^th^, 2023) and this number is likely underestimated. The use of long-reads has been particularly relevant for plant genomes due to their high repeat proportion and propensity for polyploidy. Within *Vaccinium*, high quality genomes have been assembled for nine species (Colle *et al*. 2019; Diaz-Garcia *et al*. 2021; Wu *et al*. 2021; Yu *et al*. 2021; Cui *et al*. 2022; Kawash *et al*. 2022; Yang *et al*. 2022; Mengist *et al*. 2023), as well as an ongoing pangenome project for cultivated blueberry and cranberry involving 32 cultivars (Edger 2023). In contrast, genomics of lingonberry is understudied; only a handful of genetic, chloroplast or mitochondrial genomic research has been conducted (Garkava-Gustavsson *et al*. 2005; Debnath 2007; Gailīte *et al*. 2020; Kim *et al*. 2020; Tian *et al*. 2020). This study aims to provide useful genomic resource to the lingonberry community, through genome assembly of the two distinct subspecies: *Vaccinium vitis-idaea* ssp. *vitis-idaea* and ssp. *minus*. The resources created from the study will be publicly available, in the hope of furthering our understanding of lingonberry evolution and aiding the future breeding efforts by accelerating the molecular screening of lingonberry cultivars.

## Results & Discussion

### Sequencing and assembly

Collectively, 35.3 Gbp (∼50.0X) of clean (≥ Q10) long-read data was generated (read N50 = 20.56 kbp), and additional 12.42 Gbp (∼37X) of short-read data was generated for the commercial subspecies, LC1. The *de novo* assembly resulted in 757 contigs of total length 548.004 Mbp with BUSCO (Complete) = 96.6% and contig N50 = 1.170 Mbp, and per-base accuracy = 99.959%. Similarly, 28.6 Gbp (∼46.9X) of clean long-read data (read N50 = 23.16 kbp) and 10.9 Gbp (∼35X) of short-read data were generated for the wild subspecies, LW1. The final *de novo* assembly had 518.642 Mbp of total assembly length with contig N50 = 1.400 Mbp, BUSCO (Complete) = 96.8%, and per-base accuracy = 99.975% (Table 1; Supplementary Tables 2 & 3). The assembled genome sizes were consistent or slightly smaller than flow cytometry estimates which measured a ∼550 Mbp genome (Redpath *et al*. 2022). Compared to the short-read only assemblies which generally do not reach N50 of 1 Mbp, our ONT-based assemblies are significantly more contiguous (Rhie *et al*. 2021), and our assembly statistics are comparable to many draft genome assemblies of similar size (e.g., Marrano *et al*. 2020; Wu *et al*. 2021; Hamilton *et al*. 2023; Zhang *et al*. 2023).

**Table 1:** Genome assembly statistics. Note that the haploid only assembly (for a diploid genome) means heterozygous alleles are represented as a mixed haplotype from either of the homologous copy, but not both. The allelic sequences with less confidence were purged during assembly correction based on sequence coverage (Roach *et al*. 2018).

| *V. vitis-idaea* ssp. *vitis-idaea* (LC1) | De novo assembly | Haploid only | Scaffold assembly |
| --- | --- | --- | --- |
| Total length (Mbp) | 614.857 | 548.004 | 548.071 |
| Contig N50 (Mbp) | 1.028 | 1.170 | 1.170 |
| Scaffold N50 (Mbp) | - | - | 43.867 |
| #fragments/contigs | 1358 | 757 | 757 |
| #scaffolds | - | - | 92 |
| BUSCO (C%) | 96.8 | 96.6 | 96.5 |
| BUSCO (S%) | 84.3 | 87.5 | 88.4 |
| BUSCO (D%) | 12.5 | 9.1 | 8.1 |
| QV score | 33.8254 | - | - |
| Accuracy (1-error rate) | 99.959% | - | - |
| Genome anchored to chr (%) | - | - | 98.0 |
| #genes annotated | - | - | 27,243 |
| Coding gene content (%) | - | - | 7.59 |
| TE content (%) | - | - | 45.82 |
| *V. vitis-idaea* ssp. *minus* (LW1) | De novo assembly | Haploid only | Scaffold assembly |
| Total length (Mbp) | 545.497 | 518.642 | 518.704 |
| Contig N50 (Mbp) | 1.309 | 1.400 | 1.400 |
| Scaffold N50 (Mbp) | - | - | 42.799 |
| #fragments/contigs | 1030 | 696 | 696 |
| #scaffolds | - | - | 76 |
| BUSCO (C%) | 96.9 | 96.8 | 96.9 |
| BUSCO (S%) | 89 | 89.7 | 90.5 |
| BUSCO (D%) | 7.9 | 7.1 | 6.4 |
| QV score | 35.9577 | - | - |
| Accuracy (1-error rate) | 99.975% | - | - |
| Genome anchored to chr (%) | - | - | 98.5 |
| #genes annotated | - | - | 25,718 |
| Gene content (%) | - | - | 7.37 |
| TE content (%) | - | - | 44.58 |

Scaffolding was performed by mapping to the nearest relative with a chromosome-scale genome, bilberry (*V. myrtillus*) (Schlautman *et al*. 2017; Kim *et al*. 2020; Fahrenkrog *et al*. 2022), resulting in the total of 92 and 76 scaffolds, scaffold N50 = 43.867 Mbp and 42.799 Mbp, and 98.0% and 98.5% of the contigs anchored to chromosomes for LC1 and LW1, respectively (Table 1). We characterized genomic differences between the subspecies using SyRI and found no major translocations, perhaps due to common scaffolding, but low levels of genome wide divergence in sequence (Supplementary Tables 4 & 5, Supplementary Figure 1 & 2). We recognize that reference-based scaffolding of the genome does not necessarily produce the real genome structure of lingonberry. This is because the true structural variations could be rearranged during scaffolding as the algorithm orients and places contigs based on alignment to the reference genome (Alonge *et al*. 2019). That being said, a recent study in *Eucalyptus* scaffolded ONT genomes on congeneric reference genome to study genome structure evolution and found that a very small proportion of synteny breakpoints were at contig joins, as might be expected if scaffolding is inducing false rearrangements (Ferguson *et al*. 2023). Therefore, the two lingonberry genomes created in this study can reasonably serve as a reference genome to identify genes, polymorphic genetic markers and compare with related species. Future efforts could generate an unbiased scaffolding using Hi-C or optical mapping and additionally test for the amount of bias introduced by scaffolding to a related reference genome.

### Annotation

RNA sequence data was produced from leaf sample (∼7.8 Gbp), rhizome (∼6.9 Gbp), flower (∼11.4 Gbp), and berry (∼11.7 Gbp) samples in the commercial subspecies. In addition, transcripts data from a published work was added to our analysis (∼22.5 Gbp; Tian *et al*. 2020). With the alignment of RNA reads to the genomes, the total of 27,243 and 25,718 genes were annotated (BUSCO (C): 91.4% and 91.7%). Excluding non-coding sequences (introns, untranslated regions, etc.), the coding sequence content was 7.59% and 7.37% across the genome, with the average length of 238 bp and 231 bp for LC1 and LW1, respectively. TEs were also annotated using multiple independent programs and found to cover 45.82% and 44.58% of the genome overall (Table 1). We observed that TE density was fairly even across the genome whereas genes were tended to locate less around the centre of the chromosomes and more on the chromosome arms (Figure 2). When plotting the TE distributions by different types (Supplementary Figure 3), some differences in density across the chromosome were observed.

**Figure 2:**
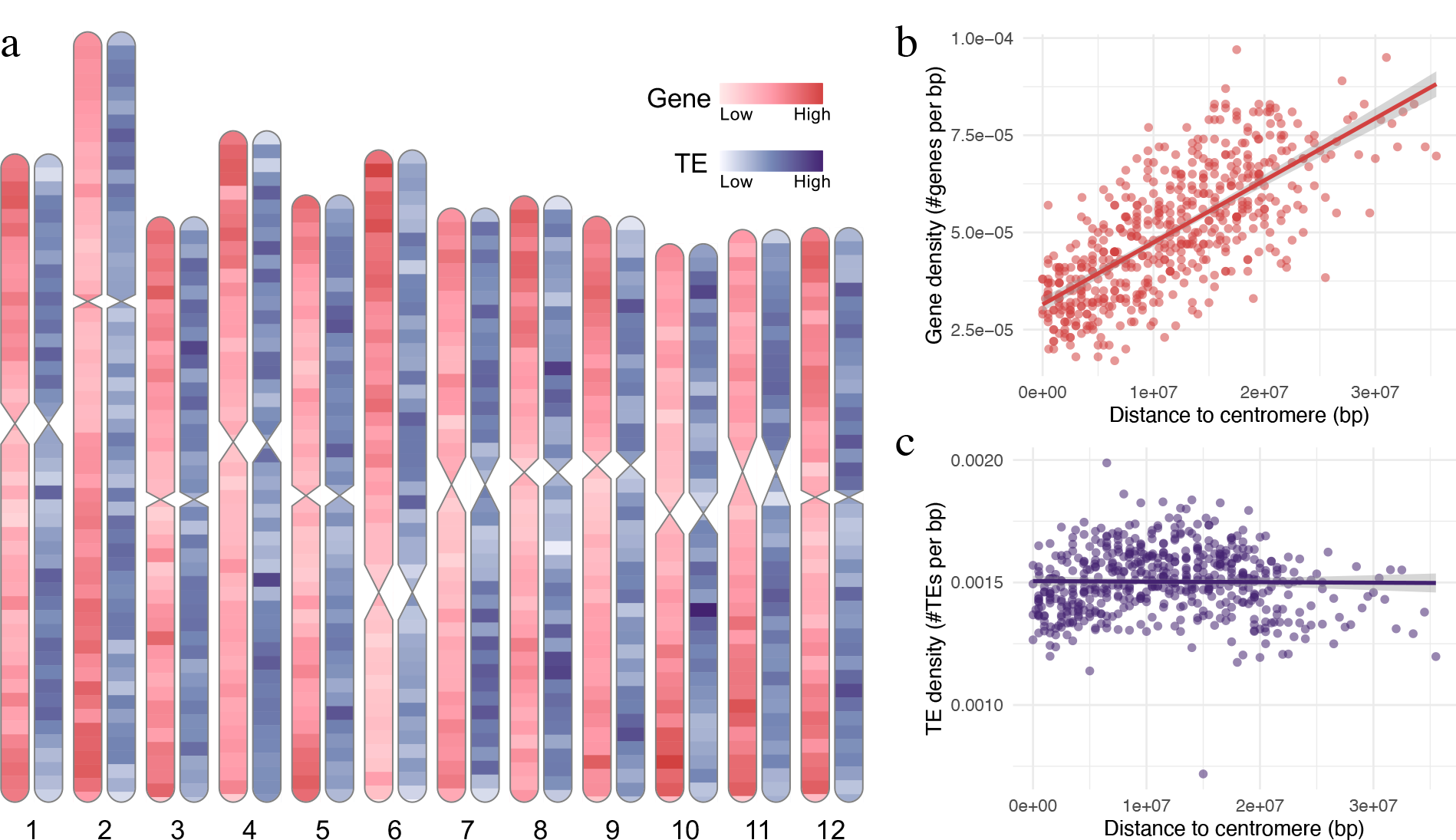
**a)** Gene and transposable element (TE) distributions in lingonberry genome (*Vaccinium vitis-idaea* var. ‘Red Candy’). **b)** Gene and **c)** TE densities by distance from centromeres. Centromere positions are approximately mapped from bilberry genome as a range, and distance was calculated to its middle value (Wu *et al*. 2021). Red shades indicate the gene density and purple shades indicate the TE density. Genes were filtered to represent only the longest gene in case of isoforms and splicing variants present. All densities are presented as the number of feature counts per 1 Mbp, except the terminal windows <1 Mbp.

**Figure 3:**
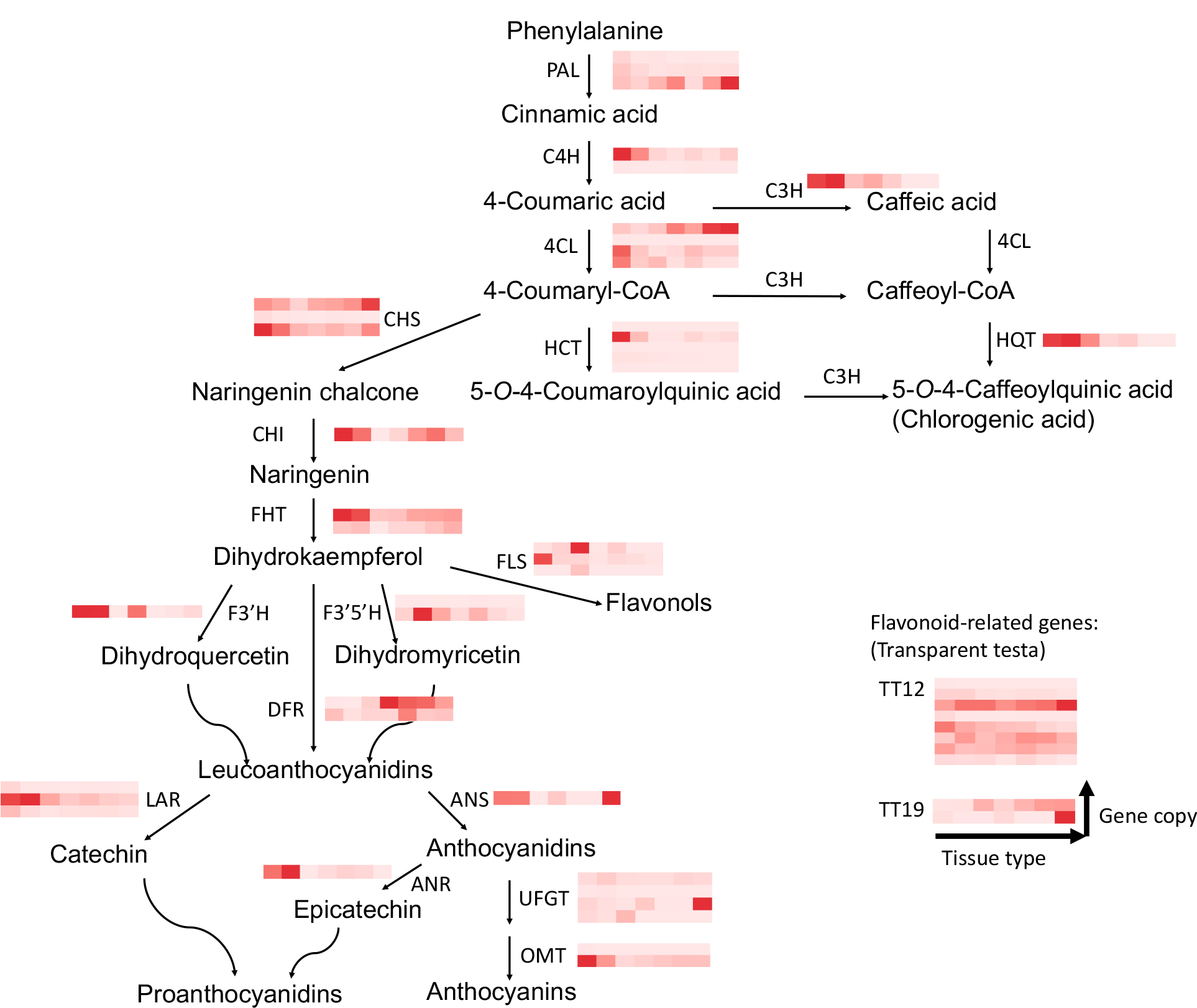
Heatmap of gene abundance related to flavonoid biosynthesis. Columns represent sample type and rows represent gene copies on lingonberry genome. Sample types are (from left to right) *Vaccinium vitis-idaea* var. ‘Red Candy’ rhizome, leaf, flower, berry, and var. ‘Sunna’ berries at different ripening stages; green berry, white berry, and red berry (Tian *et al*. 2020). Abundance is measured by fragments per kilobase of transcript per million mapped fragments (FPKM). Note that the red colour gradient is normalized within each heatmap, so comparison cannot be made across heatmaps. The enzyme pathway is based on Colle *et al*. (2019).

### The flavonoid biosynthesis pathway in lingonberry

In this study, 51 putative flavonoid pathway related genes composed of 20 distinct enzymes/structural gene categories were identified in lingonberry through orthology to the tetraploid commercial blueberry genome (*V. corymbosum* var. ‘Draper’, Colle *et al*. 2019; see Supplementary Table 6 for abbreviations and positions in lingonberry genome). Although the flavonoid related genes were expected to be highly expressed in berries compared to other tissue types, rhizome and leaf expressed C4H, HCT, HQT, CHI, FHT, F3’H, LAR, and ANS at much higher levels than the berry samples (Figure 3). Considering various known physiological roles of flavonoids in plants (Albert *et al*. 2022), abundant expression of genes in vegetative tissues implies that flavonoids play roles in stress tolerance. However, follow-up functional studies are needed to confirm their physiological roles.

While anthocyanins and the related flavonoids are the major targets of breeding due to their health benefits (Edger *et al*. 2022) and there has been efforts to build QTL maps associating genomic regions to increased anthocyanin production in commercial blueberry and cranberry (Diaz-Garcia *et al*. 2018; Montanari *et al*. 2022), the genetic basis for anthocyanin biosynthesis in lingonberry is relatively understudied. The QTL study that specifically targeted the increased anthocyanin production in blueberry suggested candidate genes including BADH acyltransferase and UFGT to be highly correlated with the increased anthocyanin profile (Montanari *et al*. 2022).

We were able to annotate four copies of UFGT in lingonberry genome, one of which was highly expressed in red berries (STRG.15162 on chromosome 4; Supplementary Figure 4). The genomic resource created in this study could be used to find such orthologues and provide a starting point to develop a set of lingonberry-specific markers which can be useful to accelerate the breeding efforts by encouraging marker-assisted selection.

**Figure 4:**
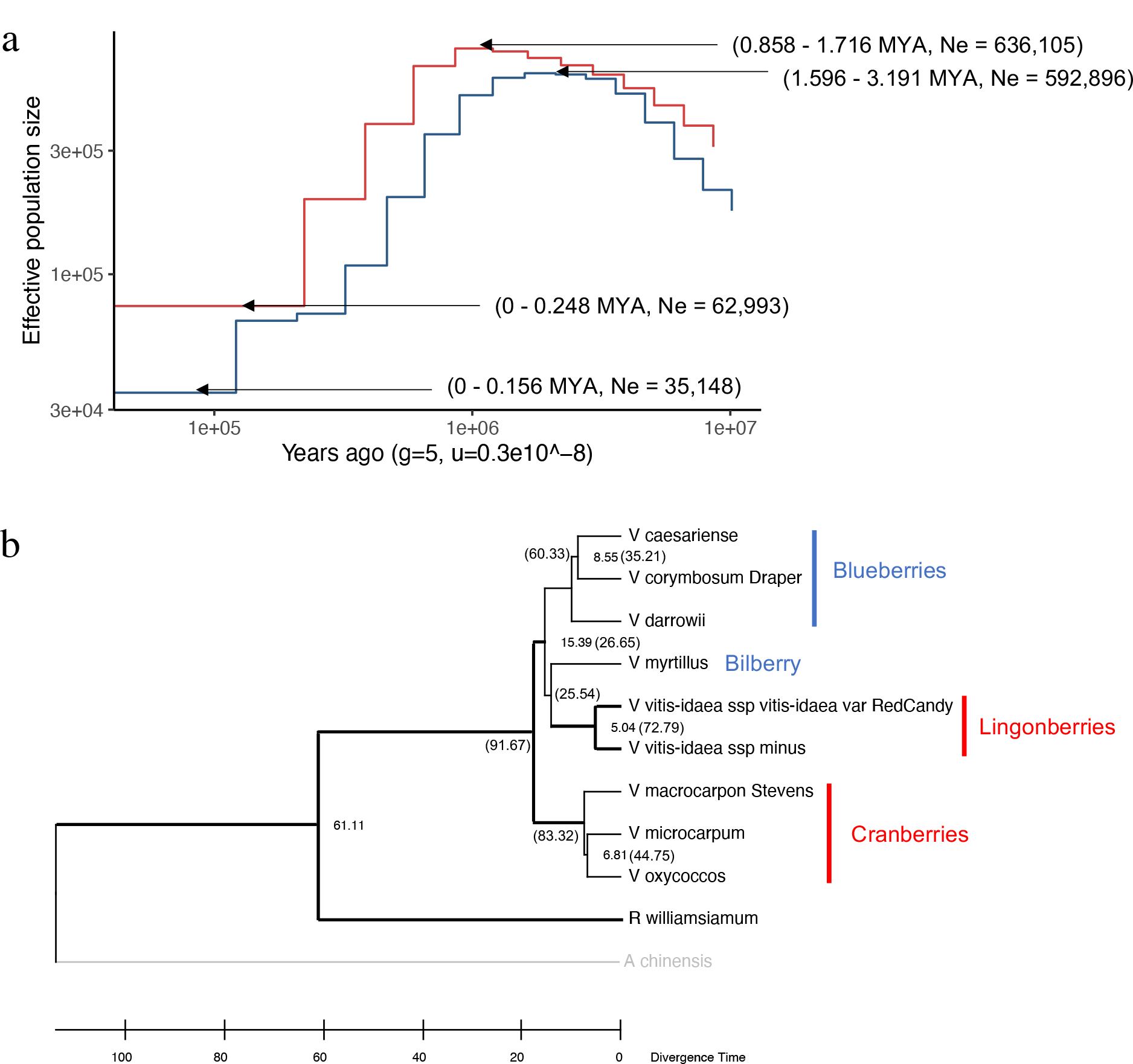
**a)** Past effective population size (Ne) of lingonberry with MSMC2. The Ne of *Vaccinium vitis-idaea* ssp. *minus* (blue; LW1) and *V. vitis-idaea* ssp. *vitis-idaea* var. ‘Red Candy’ (red; LC1) is plotted against years before present. Both x- and y-axes are log-scaled. Plots are generated with the generation time of 5 years and mutation rate of 3 x 10^9^ mutations/generation. Note the timings are presented as the range estimate from generation time of 5 to 10 years. **b)** Phylogeny of *Vaccinium* based on 2,226 conserved BUSCO genes. Thick lines indicate nodes supported by >60 STAG support values in OrthoFinder (Emms and Kelly 2018, 2019). The numbers on the selected node represents divergence time in million years (Myr), calibrated at the divergence time with *Rhododendron* (45.5–76.9 Myr), and the number in bracket shows the gene concordance factor (0–100) obtained from 2,226 BUSCO genes.

### Historical population size and origin of lingonberry

The genetic structures of the contemporary populations can often be shaped by the isolation history, which is especially relevant among sub-arctic/alpine plants that underwent past population fragmentation due to ice sheets during Pleistocene (Hewitt 2000; Eidesen *et al*. 2013).

Previous genetic studies in lingonberry revealed the impact of repeated glaciation on its contemporary patterns of genetic diversity (Debnath 2007; Eidesen *et al*. 2013). Leveraging the genome-wide variant calling along chromosomes, we were able to estimate the historical effective population size (Ne) using PSMC and MSMC2. Despite current range expansions, our result indicated an ongoing population bottleneck for both European (LC1) and North American (LW1) populations. Using a generation time of 5–10 years, we estimated that LC1 and LW1 began declining in Ne around 0.8–1.7 MYA and 1.5–3.2 MYA (Figure 4a). Lingonberry has likely undergone repetitive range contractions followed by expansion due to ice sheets advancing and receding, which may explain the population size declines over the last 1–2 MYA.

At species level, we generated a phylogeny using all the available *Vaccinium* whole-genome data. The protein sequence alignment across eight *Vaccinium* species and two outgroup species resulted in the total number of 377,681 genes analyzed, of which 349,420 were categorized into 31,264 orthogroups by OrthoFinder (Emms and Kelly 2019). The mean orthogroup size was 11.2 genes and 5,941 orthogroups were shared by all the species, of which 241 were single-copy orthogroups. Additionally, we built species tree based on 2,226 conserved single-copy BUSCO genes to confirm the congruence with the OrthoFinder result using Astral (Zhang *et al*. 2018).

We found that generally there were monophyletic groups for cranberries (*V. microcarpum, V. oxycoccos, V. macrocarpon*) and blueberries (*V. darrowii, V. caesariense, V. corymbosum*), while bilberry (*V. myrtillus*) was identified as the closest relative of lingonberry (*V. vitis-idaea*) (Figure 4b), in agreement with the previous studies (Schlautman *et al*. 2017; Kim *et al*. 2020; Fahrenkrog *et al*. 2022). Interestingly, this suggests that there are multiple colour changes of berries in *Vaccinium* lineage. Adding more species to the current tree, particularly those closely related to lingonberry and bilberry, could address whether the red berry phenotype had convergently evolved in cranberry and lingonberry lineage. Gene concordance values were generally low especially among species in the blueberry, bilberry, and lingonberry (ranging from 25–35).

Although further analysis is required to fully understand the relationship, it implies that *Vaccinium* has high levels of incomplete lineage sorting or possibly introgression between species (Coyne and Orr 2004; Beeler *et al*. 2020).

Our time calibrated phylogeny suggests that the two lingonberry subspecies diverged 5 million years (Myr), which is similar in scale to sister species divergence times in cranberry (6.8 Myr) and blueberry (8.5 Myr). We express some caution in our exact timing because this is based on a single fossil calibration and there is a lack of fossil or geological data in the younger interspecies nodes (Kumar *et al*. 2017). Compared to previous estimates, our divergence times are consistent (Cui *et al*. 2022), or overestimated (Diaz-Garcia *et al*. 2021). Nevertheless, the relative divergence between lingonberry subspecies and other *Vaccinium* species pairs suggests that the subspecies are near the divergence level expected between species and raises questions about their taxonomic classification. Further work is needed to evaluate where crossability barriers exist between ssp. *minus* and ssp. *vitis-idaea*, although high crossability is common between recognized *Vaccinium* species (Edger *et al*. 2022). The relatively old divergence time means that the parallel population bottlenecks in both subspecies are not shared, but instead independent events.

## Conclusions

This study defined the subspecies divergence in lingonberry at the whole-genome scale using genome assemblies. With the sequence variants detection and genome annotations, basic genomic knowledge was built for lingonberry including phylogenetic position in the genus and genes involved in flavonoid biosynthesis pathway. The data generated in this study will facilitate future work, such as generation of genetic markers for breeding and analysis of population structure across the species range. Further, the results encouraged future scientists in the field to address novel hypotheses regarding not only the evolution of lingonberry, but also the evolution of diverse edible berries in the genus *Vaccinium*.

## Methods & Materials

### Plant material

The clones of a commercial lingonberry plant (*Vaccinium vitis-idaea* L. ssp. *vitis-idaea* var. ‘Red Candy’) were obtained from Lochside nursery (Victoria, BC) in September 2021 and July 2022 and kept in the greenhouse, designated as LC1 and LC2, respectively. Since LC1 and LC2 are clones of the same line, they should be genetically identical, but we have used separate identifiers for each. The original plants were claimed to be collected from a wild-grown stand (location unknown). The wild lingonberry clone (*V. vitis-idaea* L. ssp. *minus*) designated as LW1, originally collected from Baie-Trinite, Quebec, Canada (Latitude: 49° 25’N; Longitude: 67° 18’W; Debnath 2007) was obtained from collaborators at Agriculture and Agri-Food Canada St. John’s Research and Development Centre, NL, and kept in the greenhouse. The three accessions were vouchered and deposited at University of Victoria herbarium collection: LC1 = UVIC 48749, LC2 = UVIC 48750, LW1 = UVIC 48751, respectively.

### High-molecular-weight DNA extraction

We excised young and mature shoots (1-2 g dry weight) from each subspecies (LC1, LW1), and wiped with 70% ethanol prior to extractions. The sterilized leaves were flash-frozen in liquid nitrogen and ground into fine powder using mortar and pestle (∼5 min). We extracted high-molecular-weight (HMW) DNA using Nucleobond^®^ HMW DNA extraction kit (Takara Bio) following the manufacturer’s protocol, with double the amount of starting material and the buffers accordingly. We then size-selected the DNA using SRE-XS kit or SRE kit (Circulomics) to remove fragments smaller than 10 kbp or 25 kbp, respectively.

### RNA extraction

We extracted the total RNA was extracted for the commercial lingonberry clones, LC1 or LC2 from five tissue types: young expanding leaf (LC1), flower (LC2), unripe berry (greenish white; LC2), ripe berry (red; LC2), and rhizome (LC2). Note that the rhizome was technically an underground shoot, but it does not have green leaves. The root-equivalent tissue could not be sampled due to soil contaminations and difficulty in extracting enough root mass without killing the plant. For leaf and flower samples, modified CTAB protocol was used to isolate RNA (Muoki *et al*. 2012; Yoshida *et al*. 2015). For rhizome, Spectrum™ Plant Total RNA Kit (Sigma) was used. For berries (LC2), modified CTAB protocol optimized for bilberry was used (Jaakola *et al*. 2001). Due to low recovery of pure RNA, the unripe and ripe berries were combined to make up one berry sample, resulting in the total of four RNA samples prepared for sequencing.

### Sequencing

For long-read sequencing with Oxford Nanopore Technologies (ONT), sequencing libraries were prepared with the Ligation Sequencing Kit (SQK-LSK110 or SQK-LSK114) and they were sequenced on MinION Flow Cell R9 (FLO-MIN106D) or R10.4.1 (FLO-MIN114), respectively, following manufacturer’s protocols. For LC1, one each of the R9 flow cell and R10.4.1 flow cell was used. For LW1, three R10.4.1 flow cells were used. All the raw output FAST5 reads were then basecalled by Guppy basecalling software v6.1.2+e0556ff (https://nanoporetech.com/) and minimap2 v2.22-r1101 (Li, 2018) using super accurate or ‘sup’ model (-c dna_r9.4.1_450bps_sup.cfg). For reads generated with R10.4.1 flow cells, the reads were further duplex-basecalled according to the Guppy Duplex-basecalling pipeline v6.3.8+d9e0f64 (https://nanoporetech.com/). In brief, raw FAST5 files were basecalled using the ‘fast’ model (dna_r10.4_e8.1_fast.cfg), and the duplex candidates were listed as read-pair candidates. Those reads were then duplex-basecalled by Guppy-duplex. The remaining reads were identified on the simplex reads already basecalled by ‘sup’ model (dna_r10.4_e8.1_sup.cfg) using a custom perl script, and finally the duplex basecalled reads were combined with the duplex-filtered simplex reads. The generated FASTQ files were concatenated as a single raw-reads output for the downstream procedures. Note that the raw basecalled reads were filtered by the mean >Q10 prior to concatenating. For short-read sequencing, the DNA library was prepared as PCR-free genome and was sequenced on Illumina NovaSeq paired-end mode, targeting 75M individual reads per sample. The RNA library was prepared by PolyA+ mRNA Library Construction service provided and sequenced on Illumina NovaSeq paired-end mode, targeting 50M reads per sample. The raw output FASTQ files were visually quality checked with fastqc v0.11.9 (Andrews 2019).

### Assembly and polishing

For LC1 assembly, initial draft genome was assembled with SmartDenovo v1.4.0 (Liu *et al*. 2021), polished with the ONT reads three times using NextPolish v1.4.0 (Hu *et al*. 2020) and with Illumina reads three times using Pilon v1.24 (Walker *et al*. 2014). In brief, the raw FASTQ paired-end reads were first filtered and trimmed using Trimmomatic v0.39 (Bolger *et al*.2014)(parameters used are ILLUMINACLIP:TruSeq3-PE.fa:2:30:10:2:True SLIDINGWINDOW:4:15 LEADING:3 TRAILING:3 MINLEN:36). The successfully paired reads were aligned to the respective long-read polished draft genome by BWA mem v0.7.17 (Li, 2013), then sorted and indexed with samtools v1.10 (Danecek *et al*. 2021) prior to polishing with Pilon for a total of three rounds with default parameters. Lastly, haplotigs and other redundant contigs were removed using purge_haplotigs v1.1.2 (parameters -l 5 -m 42 -h 95 -j 70 -s 70)(Roach *et al*. 2018). For LW1 assembly, raw ONT reads were corrected and trimmed with Canu and then assembled by SmartDenovo. The draft assembly was similarly polished with ONT reads using NextPolish three times, with Illumina reads three times using Pilon (same parameters as LC1), and haplotigs were removed using purge_haplotigs (paramters -l 5 -m 40 -h 95 -j 70 -s 70). Note that each polishing step was done up to three rounds, or until the BUSCO score started to decline. The *de novo* assembled genome was then scaffolded to chromosomes based on mapping contigs to the bilberry genome (*V. myrtillus*; Wu *et al*. 2021), using Ragtag v2.1.0 (Alonge *et al*. 2019). We did not enable the ‘correction’ mode on Ragtag, meaning it was not looking for potential misassemblies in the *de novo* assembled contigs because “misassemblies” may represent genome structure differences between bilberry and lingonberry. The final genome assembly was assessed for contiguity (N50, N90 values), per-base accuracy (QV score or consensus accuracy, error rate) and completeness (Benchmarking sets of Universal Single-Copy Orthologs; BUSCO) using BBMap v38.86 (Bushnell 2014), Merqury meryl v1.4 (Rhie *et al*. 2020) and BUSCO v5.1.2 with parameters: --lineage_dataset eudicots_odb10, --mode genome (Simão *et al*. 2015; Manni *et al*. 2021), respectively.

### Gene and TE annotation

We performed evidence-based gene annotations following the advice from the unpublished work (Freedman, *et al*. 2023 at https://github.com/harvardinformatics/GenomeAnnotation), which is particularly relevant for non-model species that lacks reliable gene models. After adapter trimming of Illumina RNA reads with Trimmomatic v0.39 with parameters same as DNA (Bolger *et al*. 2014), the quality of reads was visually checked with fastqc, making sure that there was no sequence bias or decline in read quality throughout. Additionally, published transcriptome data from *V. vitis-idaea* var. ‘Sunna’ (green, white, red berries) was added to the dataset (Tian *et al*. 2020). The reads were then aligned to the scaffolded genome including all contigs using Hisat2 v2.2.1 with default parameters (Kim *et al*. 2019). Following alignment, transcript assembly was performed using StringTie v2.1.5 with default parameters (Pertea *et al*. 2015), and the transcripts were stored as structural definition file. Gene features (i.e., untranslated regions (UTRs), exons, introns, genes, mRNAs) were then predicted on the assembled transcripts using TransDecoder v5.5.0 (Haas 2023). The longest open reading frame (ORF) prediction (command: TransDecoder.LongOrfs) was run with -S option to ensure the orientation of the paired Illumina reads. A Blastp reference library was prepared with *Arabidopsis* and *Vaccinium* known proteins from the UniProt database, to retain homologous hits on ORFs even if they do not exceed the coding likelihood scores used to filter ORF candidates in the preceding steps. We used *Arabidopsis* and *Vaccinium* protein databases because *Arabidopsis* is the most well annotated flowering plant with gene models available in eudicots, and *Vaccinium* database was the closest published protein gene models to lingonberry, in the hope to discover berry-specific genes. Finally using this information, genes were predicted (command: TransDecoder.Predict) with the parameter --retain_blastp_hits. In cases where there were isoforms (genes of same genomic position, slightly different splicing pattern) or overlapping genes (splicing variants or conflicting candidate gene models), the longest gene hit was chosen as the best candidate sequence. The completeness of the predicted genes was assessed with BUSCO with parameters: --lineage_dataset eudicots_odb10, --mode protein (Simão *et al*. 2015; Manni *et al*. 2021).

Transposable element (TE) annotation was done following the Extensive *de novo* TE annotator pipeline v2.0.0 (EDTA; Ou *et al*. 2019). In brief, candidate TEs were identified using LTR-Finder (Xu and Wang 2007; Ou and Jiang 2019), LTRharvest (Ellinghaus *et al*. 2020), LTR_retriever (Ou and Jiang 2018), TIR-Learner (Su *et al*. 2019), generic repeat finder (Shi and Liang 2019), and HelitronScanner (Xiong *et al*. 2014), followed by RepeatModeler (Flynn *et al*. 2020) to find any missed TEs due to structural-based methods. Finally, the combined repeat libraries were filtered so that coding sequences (CDS) from my transcript-based gene annotation did not get masked by repetitive regions. Additional filters to effectively remove false positives were also provided at each step of combining multiple independent programs according to EDTA pipeline (Ou *et al*. 2019). To roughly map the locations of centromeres, centromere regions of the bilberry genome (*V. myrtillus*) were transferred to my lingonberry genomes using syntenic positions (Wu *et al*. 2021) (Supplementary Table 1).

### Flavonoid biosynthesis gene expression in different tissues

Flavonoids are important berry components for both flavour and health effects. To better understand flavonoid synthesis in lingonberry, enzymes in flavonoid biosynthesis pathway in lingonberry genome were identified and then quantified using RNAseq data. Because genes that code for enzymes in anthocyanin production would be of industry and evolutionary interest, we focused our analysis on 20 enzyme-coding genes directly involved in the flavonoid biosynthesis pathway in blueberry (Colle *et al*. 2019). We first identified gene orthology between lingonberry and other *Vaccinium* species using OrthoFinder (Emms and Kelly 2019). Note that the tetraploid ‘Draper’ protein sequences were kept as a full set preserving all four haplotypes to find a potential match in lingonberry. Orthofinder places genes into orthogroups representing orthology. Any lingonberry gene found in the same orthogroup as a blueberry flavonoid biosynthesis gene was classified as a putative lingonberry flavonoid biosynthesis gene.

Using the LC1 assembly and gene annotation file produced above as a reference, expression levels of the annotated transcripts/genes were estimated by Hisat2 with -A, -G and -e option (Kim *et al*. 2019). The abundance estimate from the seven transcript dataset (i.e., LC1 leaf, LC2 rhizome/flower/berry, green/white/red berry from Tian *et al*. 2020) was reported in the units of FPKM for each dataset, corresponding to fragments per kilobase of transcript per million mapped fragments (Zhao *et al*. 2021).

### Genomic divergence between subspecies

To calculate pairwise nucleotide divergence between the two lingonberry subspecies genomes, the 12 scaffolded chromosomes were aligned using minimap2 v 2.24-r1122 (Li, 2018, 2021) with LW1 scaffolded genome as a reference and LC1 scaffolded genome as a query (default parameters: -ax ams5 --cs=long). Following data format conversions (paftools.js sam2paf | view-f maf), the alignment file was filtered to remove duplicate alignments and the pairwise divergence was calculated per 10 kbp windows using maffilter v1.3.1 (Dutheil, Gaillard, & Stukenbrock, 2014; parameters: Subset(remove_duplicates=yes, keep=no), MinBlockLength(min_length=1000), WindowSplit (preferred_size=10000, align=ragged_left), SequenceStatistics (Pairwise Divergence)). The program computes the number of basepair mismatches based on the alignment file and report this value as the divergence in % mismatch in the specified window size. Additionally, to explore the presence of structural variations and basic sequence variations, Synteny and Rearrangement Identifier v1.5 (SyRI; Goel *et al*. 2019) was used on the aligned chromosomes with default parameters.

### Demographic history estimate

In order to investigate the past population history of lingonberry subspecies, we utilized multiple sequentially Markovian coalescent model (MSMC2; Schiffels and Wang 2020) and pairwise sequentially Markovian coalescent model (PSMC; Li and Durbin 2011). MSMC2 requires that the analyzed populations are mapped to the same reference genome. For the purpose of comparing the two methods in parallel, we chose to use LW1 as a reference genome for both subspecies because of better contiguity and basepair accuracy than LC1. To first calculate the effective population size (Ne) of each subspecies, the paired Illumina reads were mapped to the LW1 genome using BWA mem v0.7.17 (Li, 2013). PCR and optical duplicates were then removed using GATK Picard v2.23.2 ‘MarkDuplicates’ function (Van der Auwera and O’Connor 2020). The mappable heterozygous variant sites were identified separately for each chromosome per subspecies following bamCaller.py in MSMC2 v2.1.3 (Schiffels and Wang 2020). In brief, SNPs were first called using bcftools v1.16 (Danecek *et al*. 2021) with the command ‘mpileup’ and ‘call’ with the parameters: -q 20 -Q 20 -C 50 and -c -V indels, respectively. The results were then filtered and organized based on read coverage (mean coverage set to 38 for LW1, 37 for LC1; filtering applied is the minimum of x1/2 mean coverage to the maximum of x2 mean coverage). An additional mappability mask was generated to avoid calling variants from significantly repetitive regions using GenMap v1.3.0 (Pockrandt *et al*. 2020) with the parameter:-K 30 -E 2. For PSMC inputs, SNPs were similarly called using bcftools ‘mpileup’ and ‘call’ with the same parameters as above, and the results were filtered with the minimum of x1/3 and maximum of x2 mean coverage, as recommended (Li and Durbin 2011). No repeats mappability mask was considered in PSMC analysis. When running the models, a generation time of 5–10 years was chosen based on a prior experiment observing minimum of 8 years required to consider a seedling fully reproductive (Hjalmarsson and Ortiz 1998) and considering the woody shrub’s natural age of first flowering (Ritchie 1955). However, given the potential for reproduction after first maturity, we recognize that this may underestimate the average reproductive age of the natural population. A mutation rate of 3 x 10^9^ substitutions per generation from *Arabidopsis thaliana* was used (Exposito-Alonso *et al*. 2018).

### Phylogenetic tree construction

Phylogenetic trees were constructed using two different approaches. The first approach follows the default pipeline provided using OrthoFinder v2.5.4 (Emms & Kelly, 2019). In brief, a total of 11 species protein sequences in amino acid fasta format were collected from published studies: eight *Vaccinium* species: two *Vaccinium vitis-idaea* subspecies from this study, *V. corymbosum* var. ‘Draper’ v1.0 first 12 chromosomes (Colle *et al*. 2019), *V. macrocarpon* var. ‘Stevens’ v1.0, *V. microcarpum* v1 (Diaz-Garcia *et al*. 2021), *V. oxycoccos* NJ96-20 v1 (Kawash *et al*. 2022), *V. myrtillus* NK2018_v1 (Wu *et al*. 2021), *V. darrowii* v1.2 (Cui *et al*. 2022), and *V. caesariense* W85-20 P0 v2 (Mengist *et al*. 2023). Kiwi fruit or *Actinidia chinensis* v3.0 (Tang *et al*. 2019) and Azalea or *Rhododendron williamsianum* (Soza *et al*. 2019) were used as outgroups. The species tree was constructed based on the individual gene trees inferred from the orthologous gene groups as per OrthoFinder pipeline (Emms and Kelly 2017, 2018). For further validation using conserved genes only, single-copy BUSCO genes were extracted and aligned to infer species tree. To do that, BUSCO analysis was first performed on the collected genome assembly itself in nucleotide fasta format with --lineage_dataset eudicots_odb10, --mode genome (Simão *et al*. 2015; Manni *et al*. 2021). Then the identified single-copy genes were aligned by MAFFT v7.310 and the individual gene trees were inferred with IQ-TREE v1.5.5 (Nguyen *et al*. 2015). Outlier long branches were trimmed by TreeShrink v1.3.9 (Mai and Mirarab 2018) with default parameters. Finally, the species tree was constructed using the trimmed gene tress in Astral III v5.7.8 (Zhang *et al*. 2018). For visualization and data interpretation, both species trees were exported in Newick format, and then viewed in FigTree. Trees were rooted manually to *Actinidia chinensis*.

Additionally, divergence times were estimated following (Diaz-Garcia *et al*. 2021). In brief, single-copy BUSCO gene alignments were used as an input alignment file with RelTime as implemented in MEGA X (Tamura *et al*. 2012, 2018). *Actinidia chinensis* was set as an outgroup and the following calibration time was used based on the average of 16 studies in TimeTree (Kumar *et al*. 2017): *Rhododendron* and *Vaccinium* (45.5–76.9 MYA). Uniform distribution was selected as the calibration density. Due to MEGA X requiring a single sequence alignment file with equal sequence length, only 743 BUSCO genes that were present in all 11 species were selected for analysis. The individually aligned BUSCO genes were concatenated to prepare the input file with seqkit concat function (Shen *et al*. 2016). Note that seven ambiguous amino acid “J”s corresponding to isoleucine or leucine in the alignment file were manually replaced with “I”s in order to meet the requirements by MEGA X.

## Data availability statement

This Whole Genome Shotgun project has been deposited at DDBJ/ENA/GenBank under the accession JAUYVE000000000 (LC1) and JAUYVF000000000 (LW1). The raw sequences are archived in SRR25468432-50 (ONT) and SRR25477285-90 (Illumina). Full annotations and reference-mapped genome assemblies used in this manuscript can be downloaded from figshare: https://figshare.com/projects/Unveiling_the_evolutionary_history_of_lingonberry_Vaccinium_vitis-idaea_L_through_genome_sequencing_and_assembly_of_European_and_North_American_subspecies/175089. All codes used for assembly pipeline and downstream analysis are available at https://github.com/kaede0e/lingonberry_genomics.

## Author contributions

KH compiled data, assembled genome, conducted annotation and subsequent analyses. GLO supervised the project. KH and GLO conceived of the project and wrote the manuscript. SCD provided plant material, advised, and approved the manuscript.

## Acknowledgement

The authors acknowledge the collaborators at Agri-Foods Canada St. John’s Research and Development Centre for collecting, maintaining, and providing the wild lingonberry samples, and the Digital Research Alliance of Canada for computational resource and technical support. The authors acknowledge the Michael Smith Genome Sciences Centre at UBC for sequencing. The authors also thank Dr. Ben Koop, Dr. Kris Christensen, and Anne-Marie Flores for sharing their facilities and expertise throughout the project.

