## Supplementary Figure for "Unveiling the evolutionary history of lingonberry (*Vaccinium vitis-idaea* L.) through genome sequencing and assembly of European and North American subspecies"

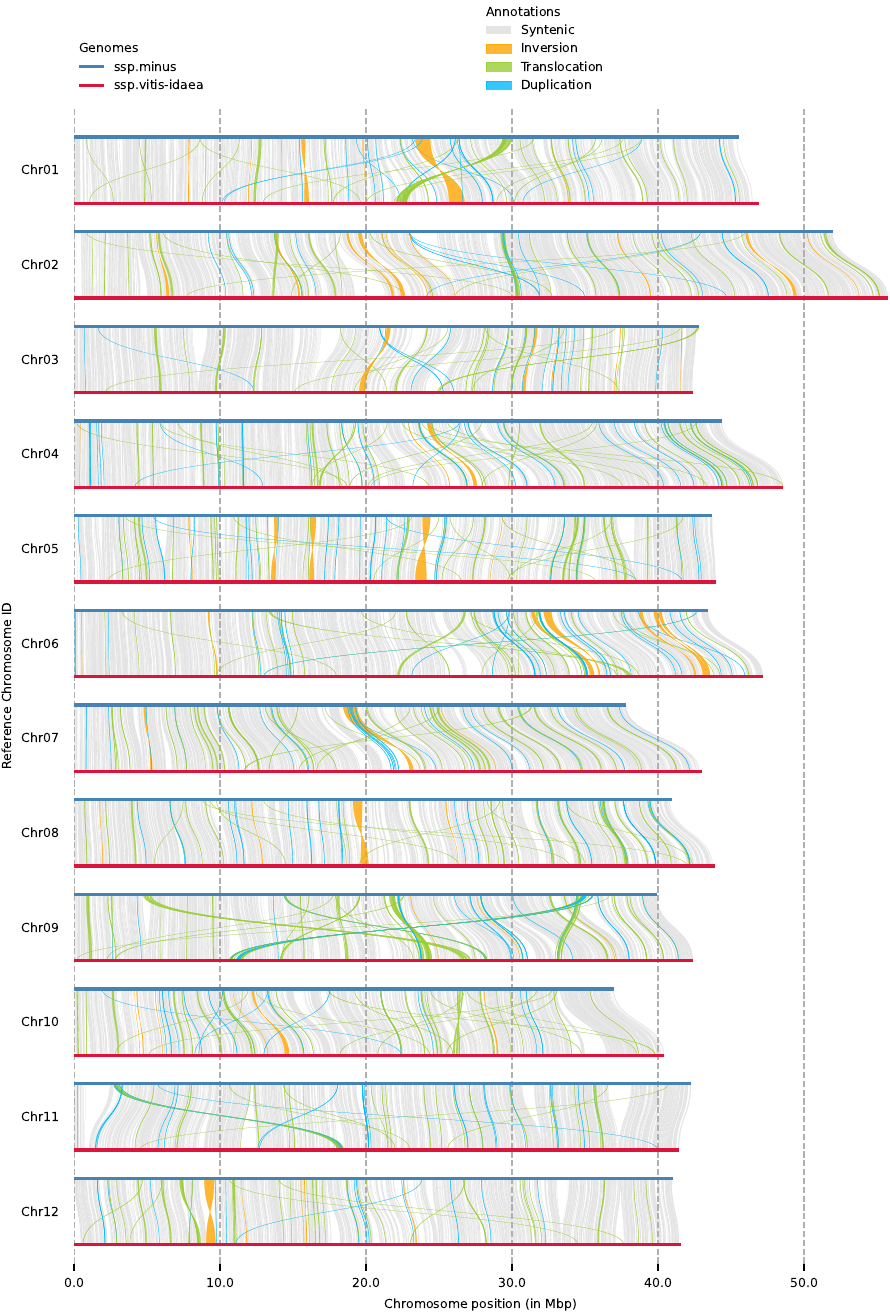

Supplementary Figure 1: Alignment between two lingonberry subspecies detected by SyRI (Goel *et al.* 2019). Horizontal lines represent chromosomes on each genome: blue is *V. vitis-idaea* ssp. *minus* (LW1), red is *V. vitis-idaea* ssp. *vitis-idaea* var. ‘Red Candy’ (LC1). Structural variations are shaded in colours: grey for syntenic region; orange, inversion; green, translocation; light blue, duplication. Plots are made with plotsr (Goel and Schneeberger 2022).

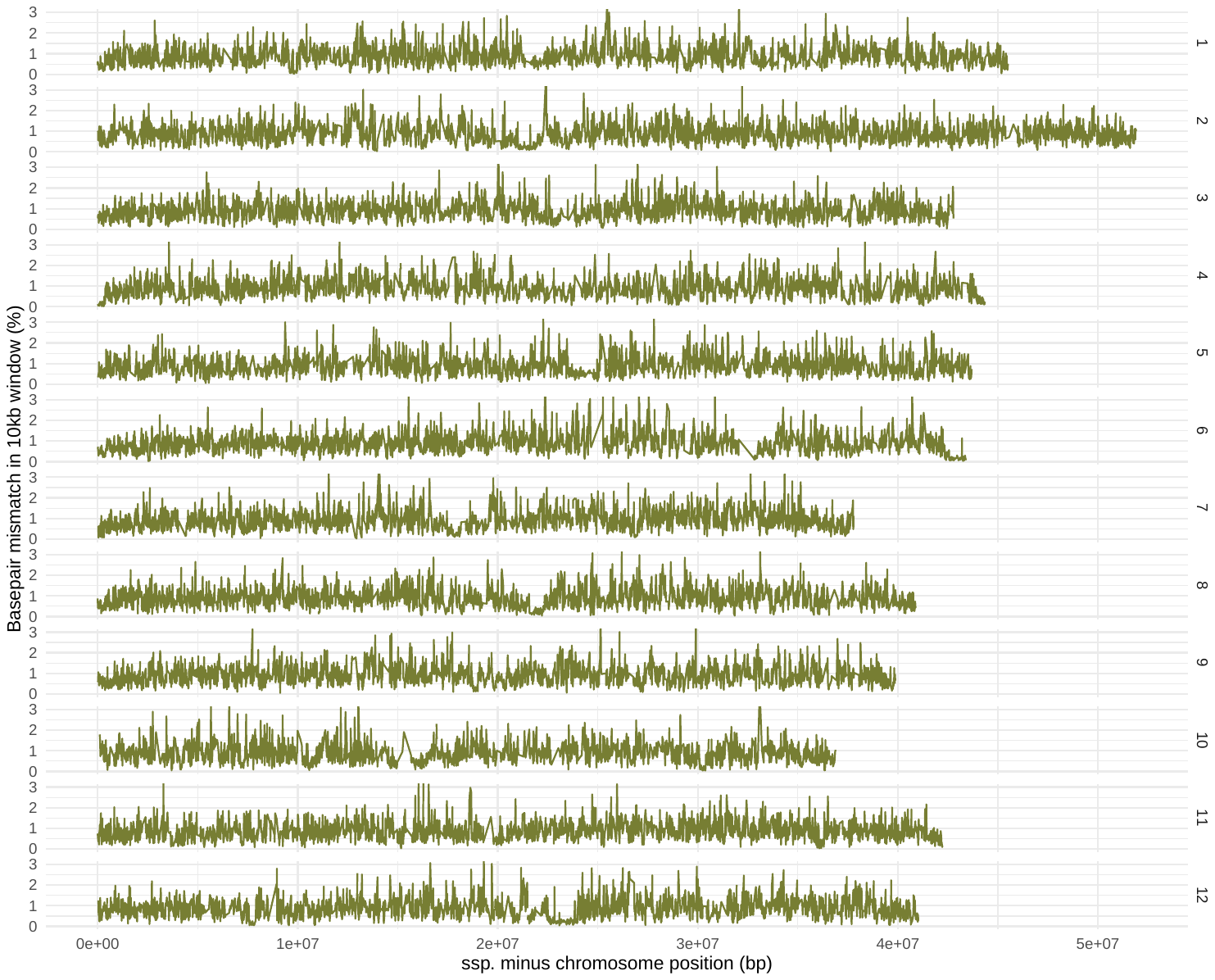

Supplementary Figure 2: Pairwise divergence between *Vaccinium vitis-idaea* ssp. *minus* (LW1) and ssp. *vitis-idaea* (LC1). The percentage of basepair mismatch was determined based on the aligned sequences in 10 kbp windows. Any alignments <1000 bp was filtered out before plotting. The reference genome was set to LW1. Rows are sorted by chromosomes as labelled on the right.

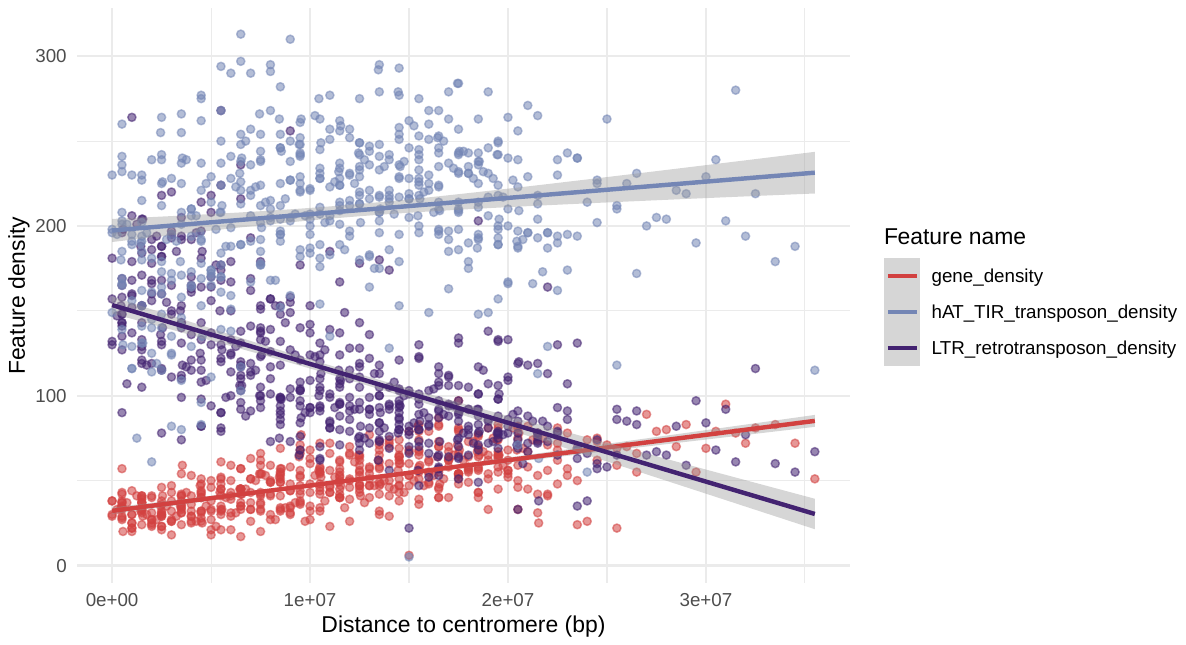

Supplementary Figure 3: Scatter plots of long-terminal-repeat (LTR) and hAT terminal inverted repeat (hAT TIR) densities in comparison to gene density against the distance from the centre of chromosomes. Red points indicate gene density, light blue indicates hAT-TIR transposon density, and purple indicates LTR retrotransposon density. All densities are presented as the number of feature counts per 1 Mbp window.

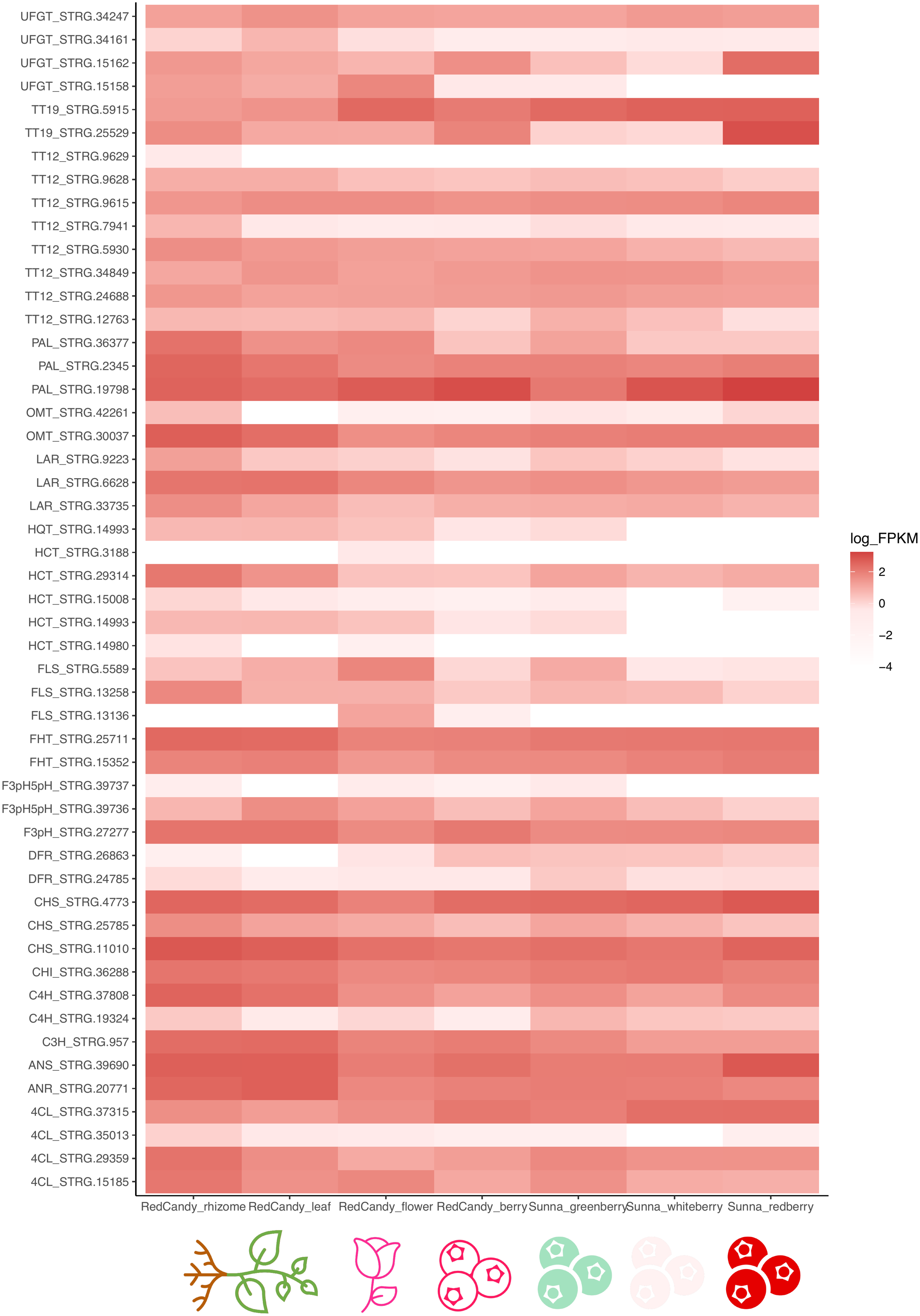

Supplementary Figure 4: Heatmap of flavonoid biosynthesis related gene abundance in lingonberry. Gene abundance was measured in the unit of FPKM, and the values are log scaled for visualization purpose. Rows represent different copies of each orthologous gene in lingonberry (enzyme name_STRG-id), and columns are sample types.
